# Transferable spatial omics deconvolution with SpaRank

**DOI:** 10.64898/2026.05.09.723936

**Authors:** Xuhua Yan, Ruiqing Zheng, Jinmiao Chen, Min Li, Wei Lan

## Abstract

By resolving cell-type compositions from multi-cellular spatial measurements, deconvolution is central to resolving the cellular landscape of complex tissues. Existing deconvolution methods fit continuous expression values and are therefore sensitive to batch effects between single-cell references and spatial data, requiring retraining for each new context. Here we present SpaRank, a context-aware framework that performs spatial deconvolution by representing spots as ranked feature sequences. Adapting the rank-based encodings of single-cell foundation models, this formulation is inherently robust to technical variation, enabling a pretrain-transfer paradigm. On simulated benchmarks, SpaRank achieves strong deconvolution accuracy, robustness to expression perturbations, and substantial computational efficiency. On experimental datasets, pretrained models generalize across diverse biological contexts: a model pretrained on a multi-organ lymphoid atlas accurately resolved cell-type distributions across distinct tissues and sequencing platforms; likewise, a model pretrained on an integrated breast atlas delineated cell-type compositions across normal and malignant disease states. Furthermore, the framework naturally extends to multimodal spatial deconvolution by employing gated fusion to adaptively integrate diverse omics signals, improving accuracy over single-modality approaches. Overall, SpaRank establishes a transferable deconvolution paradigm, enabling unified cellular atlases to support direct, context-aware inference across diverse biological states and profiling modalities.

## Introduction

Spatial omics technologies have transformed our understanding of tissue biology by mapping molecular profiles to their native histological locations^1,2^. However, widely used spatial profiling platforms often achieve transcriptome-scale coverage at the expense of spatial resolution^3,4^. As a result, each spot typically captures mixed signals from multiple cells. Computational deconvolution has therefore become an essential analytical step for resolving these mixtures. By leveraging single-cell references, deconvolution methods estimate the cell-type composition of each spatial spot. This increased resolution is critical for delineating complex tissue microenvironments, localizing disease-associated niches, and deciphering local cell-cell communication^5^. Yet, the computational deconvolution of spatial mixtures is severely limited by technical variation between single-cell and spatial platforms, as well as differences in biological context between the reference atlas and the target tissue.

At present, a variety of deconvolution algorithms have been developed, encompassing probabilistic and deep generative frameworks (e.g., Cell2location^6^, Stereoscope^7^, DestVI^8^), factorization-based regression models (e.g., CARD^9^, RCTD^10^, SPOTlight^11^), and deep learning or global mapping strategies (e.g., SPACEL^12^, Tangram^13^). These methods have demonstrated strong performance and broad applicability^14^. However, as these methods model cellular composition from quantitative expression values, they are susceptible to technical variations between single-cell references and spatial data. This sensitivity limits transferability, requiring retraining for each new dataset or tissue condition rather than direct deployment. It also hinders the use of large cellular atlases as unified references for deconvolution across diverse physiological and pathological states, eliminating the need for dataset-specific reference curation. Moreover, their reliance on fixed distributional assumptions over continuous expression values limits their extension to multimodal spatial profiling.

To address these limitations, we developed SpaRank, a transferable spatial deconvolution framework that represents each spot as a ranked gene sequence rather than a quantitative expression vector. Because relative expression rankings are more stable than absolute expression values across platforms, this rank-based formulation is inherently robust to technical variation, as supported by recent single-cell foundation models using similar representations^15,16^. This enables a pretrain-transfer paradigm in which a single model trained on a large multi-context cellular atlas can generalize across tissues, disease states, and platforms without dataset-specific retraining. Furthermore, because our framework makes no distributional assumptions about expression values, it naturally extends to spatial multi-omics deconvolution. We benchmarked SpaRank against existing methods using sections simulated from multiplexed error-robust fluorescence in situ hybridization (MERFISH)^17^ and Xenium data, demonstrating superior deconvolution accuracy. We further validated model transferability by pretraining on two distinct reference atlases—one spanning multiple organs, and one integrating normal and malignant breast profiles from independent cohorts—and directly deploying across novel spatial sections without retraining. Finally, we assessed its multi-modal capacity through simulated spatial multi-omics deconvolution benchmarks, where SpaRank outperformed existing single-modality methods, and further confirmed its applicability on real multi-modal spatial sections.

## Results

### The SpaRank model

SpaRank is a sequence-based deep learning framework for transferable spatial-omics deconvolution (**Fig. 1a**). The model is pretrained on synthetic spatial spots simulated from single-cell references, where cells sharing the same biological context (e.g., organ, disease state, or technical batch) are aggregated to approximate spot-level mixed signals, with cell-type proportions as prediction targets. For each spot, marker features derived from the reference atlas are ranked by expression level, and the top *K*_*g*_ feature names are extracted as an input sequence alongside a categorical context label. Inspired by single-cell foundation models^15,16^, SpaRank encodes this sequence via a Transformer^18^ encoder and fuses it with a context embedding to predict cell-type proportions through a regression head.

**Fig. 1.**
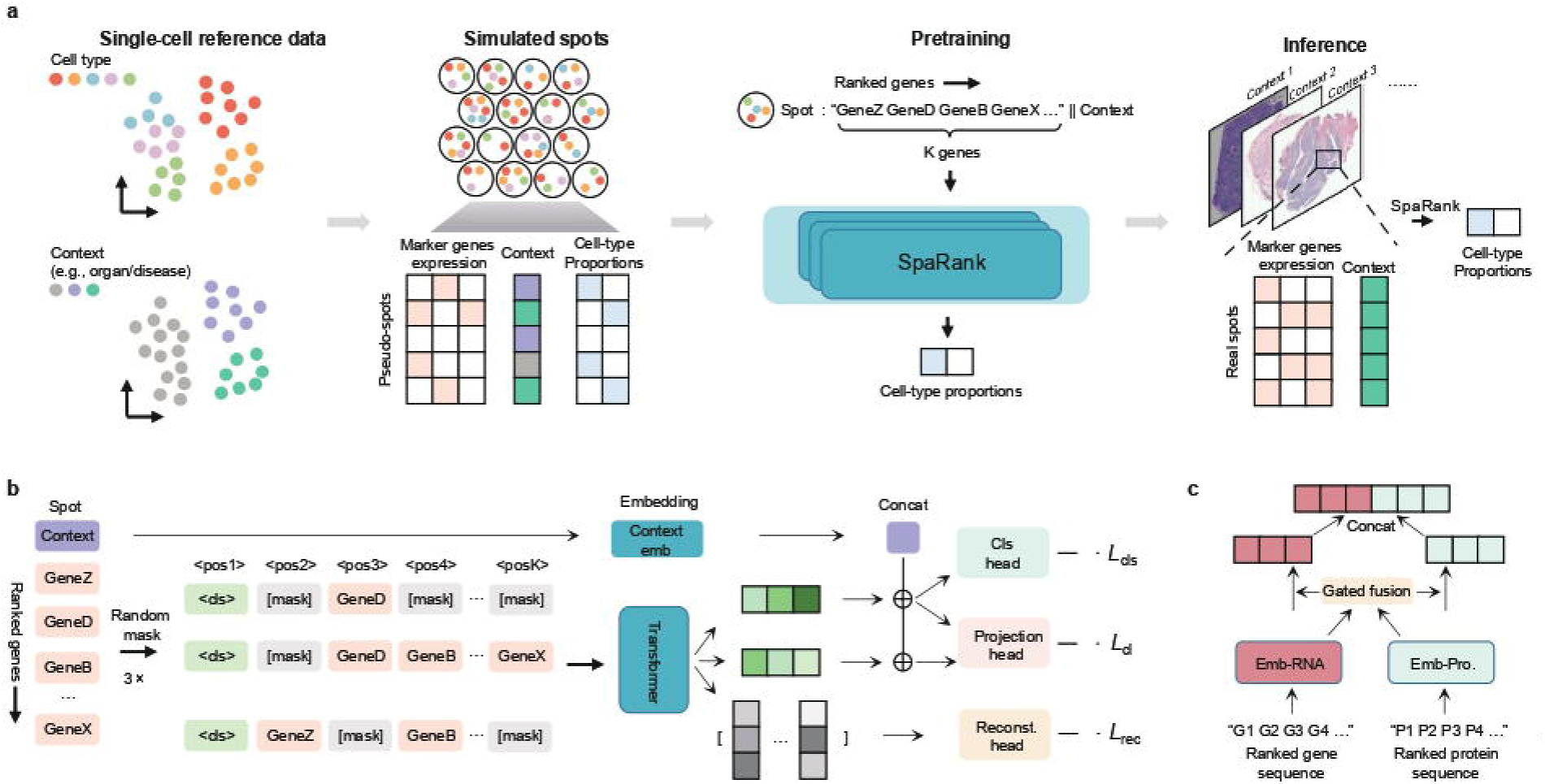
Overview of SpaRank framework. **a**, SpaRank workflow. Cells from a single-cell reference are sampled and aggregated within each biological context to generate pseudo-spots with ground-truth cell-type proportions. Expression profiles are converted into ranked gene sequences, paired with context labels, and used for model pretraining. The pretrained model is then applied directly to spatial sections from different contexts without retraining. **b**, Model architecture and training objectives. Each input sequence undergoes three independent random masking operations. The first two masked sequences a.re encoded and compared via a contrastive loss. The third is passed to a reconstruction head for masked token prediction. The first masked sequence is concatenated with the context embedding and passed to a regression head for cell-type proportion prediction. **c**, The core of multimodal extension. Each modality is encoded by a modality-specific Transformer, and a gated multimodal fusion mechanism adaptively integrates the modality embeddings.

A key challenge in training SpaRank is overfitting to the top-ranked features, which carry strong cell-type signals but may cause the model to neglect informative patterns across the full sequence. To mitigate this, SpaRank applies three independent random masking operations to each input sequence (**Fig. 1b**). Two masked variants are encoded, fused with the context embedding, and then compared via a contrastive loss^19^, forcing the encoder to remain consistent when top-ranked features are randomly perturbed or removed. The third is processed through a reconstruction head that predicts masked feature identities, compelling the model to capture signals distributed across the full sequence. The embedding from the first masked variant is then fused with the context embedding and passed to a regression head, where a cross-entropy loss is computed against ground-truth cell-type proportions.

Since SpaRank operates on ranked sequences without distributional assumptions over expression values, it generalizes naturally to multimodal spatial deconvolution. In this formulation, each modality is independently encoded by a modality-specific Transformer, integrated via gated multimodal fusion, and combined with the context embedding before the regression head yields final proportion predictions (**Fig. 1c**).

### Benchmarking on simulated mouse isocortex data

We first benchmarked SpaRank on simulated spatial data derived from a single-cell-resolution MERFISH atlas of the adult mouse brain^20^ (**Fig. 2a**), using a matched whole-brain scRNA-seq atlas^21^ as the deconvolution reference. Both datasets were restricted to the isocortex region, comprising 24 cell types. Pronounced batch effects were observed between the MERFISH and scRNA-seq data (**Fig. 2b**). The scRNA-seq reference was then downsampled to 46,575 cells across 32,285 genes, and 27 MERFISH isocortex sections were gridded into pseudo-spots with known cell-type proportions (**Fig. 2a; Methods**).

**Fig. 2.**
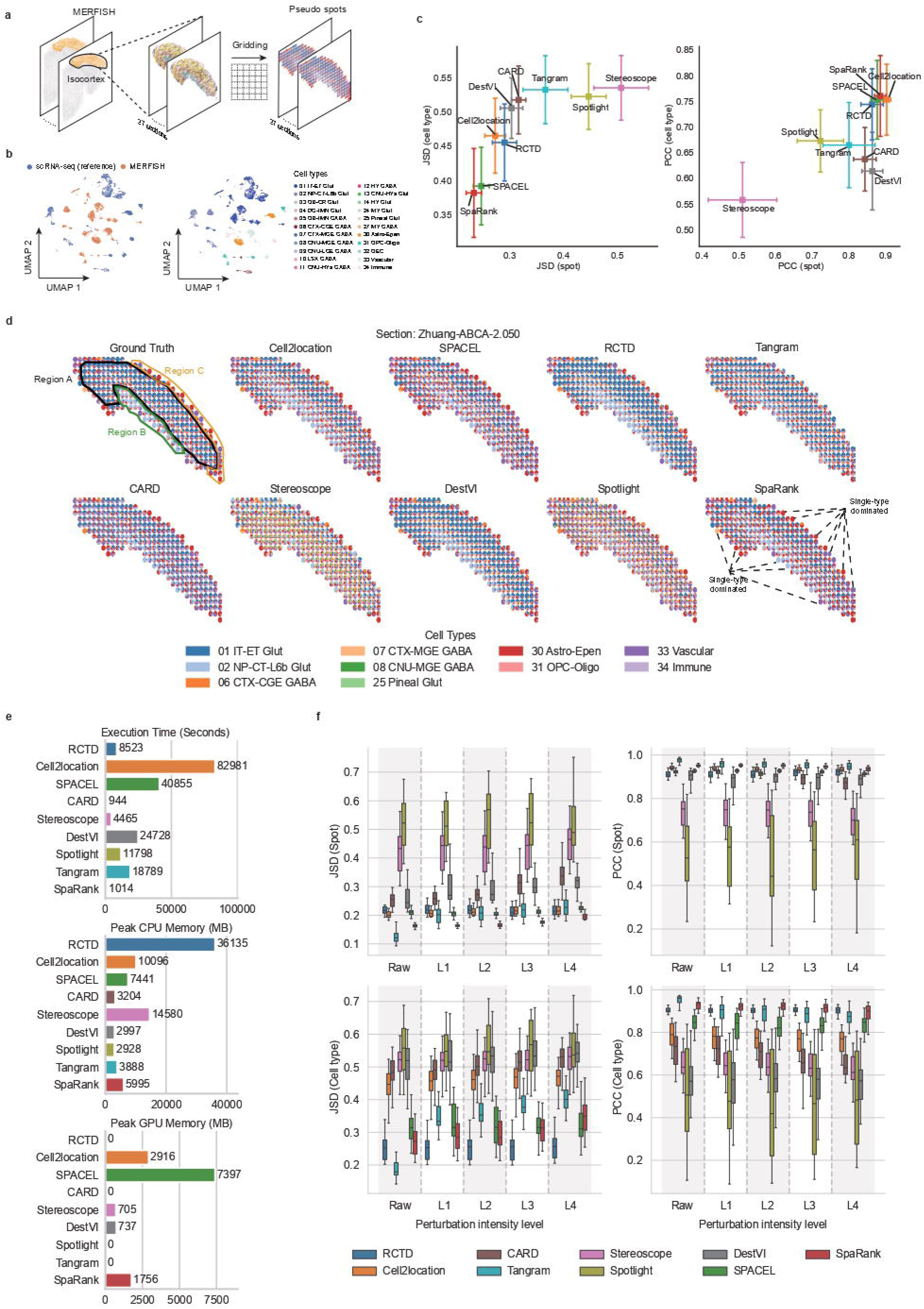
Benchmarking on simulated mouse isocortex datasets. **a**, Simulation workflow. Isocortex regions were extracted from 27 single-cell-resolution MERFISH sections and gridded into spot-resolution pseudo-sections. **b**, UMAP visualization showing pronounced batch effects between the scRNA-seq reference and MERFISH data. **c**, Deconvolution performance across 27 sections, measured by JSD (left) and PCC (right) at the spot level (x-axis) and cell-type level (y-axis). Points indicate the mean across sections for each metric; error bars indicate the standard deviation along axes. **d**, Spatial pie plots showing inferred cell-type compositions from each method on an example section. Ground-truth proportions delineate three distinct regions (A, B, C). Correctly predicted single-cell-type-dominated spots are highlighted for SpaRank. **e**, Total computational cost across all 27 sections per method, shown as bar plots for CPU time, peak CPU memory, and peak GPU memory. **f**, Boxplots of metrics across all methods under increasing perturbation of reference expression profiles. Each box summarizes results across 27 sections for a given method and perturbation intensity, with five intensity levels shown from left to right. ‘Raw’ denoting the unperturbed reference. Center line, median; box limits, upper and lower quartiles; whiskers, I.5x interquartile range.

SpaRank was compared against seven established methods: Cell2location, CARD, DestVI, RCTD, SPOTlight, Stereoscope, SPACEL, and Tangram. Among these, Cell2location, DestVI, and Stereoscope first learn cell-type signatures from the reference and then retrain on each spatial section, while the remaining methods are trained jointly on each reference-section pair. SpaRank was pretrained once on the reference and deployed directly across all 27 sections. Performance was assessed along two axes: spot level and cell-type level, using Pearson correlation coefficient (PCC) and Jensen-Shannon divergence (JSD)^22^, where higher PCC and lower JSD indicate better deconvolution accuracy.

SpaRank achieved the highest mean scores across all 27 sections on both spot-level and cell-type-level JSD, followed by SPACEL (**Fig. 2c**). Under PCC metrics, SpaRank ranked first on cell-type-level PCC, followed closely by SPACEL and Cell2location, while Cell2location led on spot-level PCC with SpaRank and SPACEL close behind. Stereoscope consistently ranked last across all four metrics. Among methods requiring section-specific retraining, Cell2location emerged as the top performer. However, SpaRank outperformed all such methods while requiring only a single pretraining step on the reference data. This result demonstrates that SpaRank’s pretrain-transfer paradigm supports robust deconvolution generalization even in the presence of pronounced batch effects between reference and target data.

Spatial maps of predicted cell-type proportions on an example section further illustrate these differences. Specifically, the ground truth revealed three distinct domains (**Fig. 2d**): a central domain dominated by IT-ET glutamatergic neurons (region A), a lower domain enriched for NP-CT-L6b glutamatergic neurons (region B), and a peripheral domain characterized by Astro-Epen cells (region C). SpaRank’s predictions closely recapitulated this spatial organization, correctly identifying single-cell-type-dominated boundary spots where competing methods introduced spurious contributions from other classes. RCTD and DestVI over-predicted IT-ET glutamatergic neurons in region A, while Stereoscope and SPOTlight assigned substantial proportions of cell types entirely absent from regions A and B, such as CNU-MGE GABA and Pineal Glut populations, consistent with their poor quantitative performance.

By eliminating the need for section-specific retraining, SpaRank conferred substantial computational advantages. Across all 27 sections, SpaRank and CARD achieved the lowest total runtime (~1,000 seconds), while Stereoscope required at least 4,500 seconds and Cell2location exceeded 80,000 seconds (**Fig. 2e**). SpaRank additionally maintained comparatively low CPU memory and GPU memory consumption.

Finally, we evaluated SpaRank’s robustness by introducing controlled perturbations into reference expression profiles to simulate varying degrees of technical variation between the reference and spatial data (**Methods**). These perturbations progressively disrupted the relative ranking structure of the reference profiles. For this robustness analysis, each simulated spatial section was paired with single-cell data derived from the same MERFISH section as the reference, providing an ideal, unperturbed reference for each target. Without any perturbation (raw), Tangram ranked first and SpaRank second across all four metrics (**Fig. 2f**). This is expected because the task in this setting is inherently equivalent to mapping single cells back to the spots they compose, which aligns precisely with Tangram’s optimization objective^13^. As perturbation intensity increased, however, Tangram’s performance degraded markedly across all metrics except spot-level PCC. SpaRank, by contrast, maintained top rankings on spot-level JSD and cell-type-level PCC under increasing perturbation, ranked second on spot-level PCC behind Tangram, and remained competitive on cell-type-level JSD. These results demonstrate that SpaRank maintains robust performance under increasing expression perturbations, supporting reliable pretrain-transfer deployment across real-world cross-dataset settings.

### SpaRank enables transferable deconvolution across diverse lymphoid tissues

To evaluate SpaRank in a realistic transfer setting, we collected a multi-organ reference atlas^6^ comprising 73,260 cells across 34 cell types from human lymph node, tonsil, and spleen^23-25^ — three related lymphoid organs with pronounced batch effects and distinct cellular compositions (**Fig. 3a; Methods**). SpaRank was trained on this unified atlas and applied, without retraining, to four spatial sections spanning different tissues and platforms: a CytAssist Visium tonsil section^26^, a CytAssist Visium lymph node section^26^, a Visium lymph node section^27^, and an Array-seq spleen section^28^ (**Methods**). All baseline methods were evaluated using the same reference atlas rather than organ-matched subsets. All sections except the Visium lymph node carried manual anatomical annotations from the original studies, providing an external reference for assessing predicted cell-type localization.

**Fig. 3.**
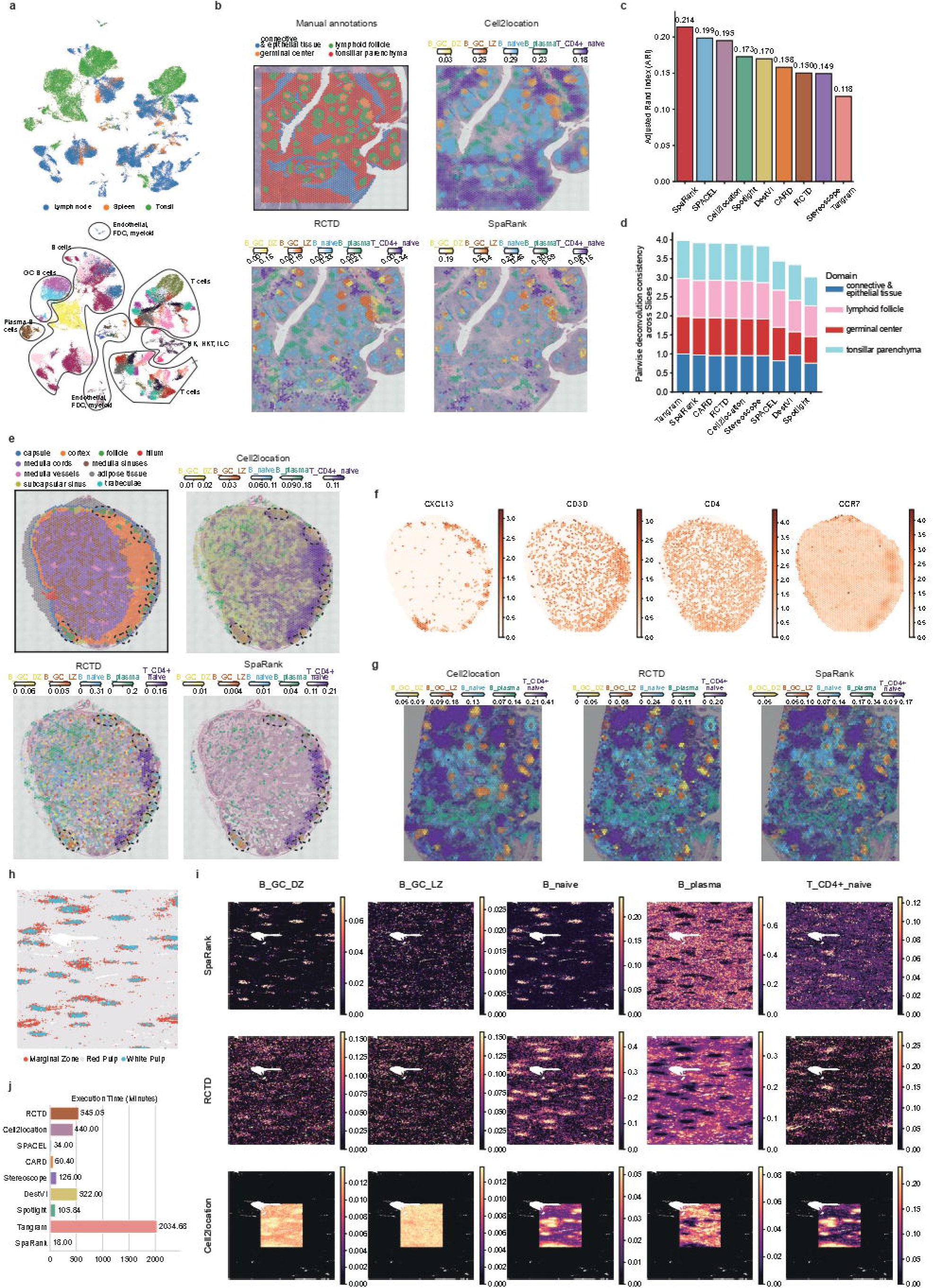
Multi-organ transferability across tissues and sequencing platforms. **a**, UMAP visualization of the single-cell reference atlas (before integration) combining data from human lymph node, tonsil, and spleen, colored by tissue origin (upper) and cell type (lower). **b**, Manual anatomical annotations and estimated cell-type abundances from SpaRank, Cell2location, and RCID on a CytAssist Visium tonsil section, shown as color intensity over the H&E image. **c**, Adjusted Rand Index (ARI) comparing clustering of predicted proportions against manual annotations for all methods. **d**,Stacked bar plot of pairwise Pearson correlations of predicted proportions across annotated domains between tonsil section replicates for all methods. **e**, Manual anatomical annotations and estimated cell-type abundances from three methods on a CytAssist Visium lymph node section, shown over the H&E image. **f**, Spatial expression maps of RNA markers *CXCLJ3, CD3D*, and *CD4*, and protein expression of CCR7. **g**, Estimated cell-type abundances from three methods on a standard Visium lymph node section, shown over the H&E image. **h**, Manual anatomical annotation of the Array-seq human spleen section. **i**, Estimated abundances of selected cell types from three methods. Cell2location results are shown on the subsampled spot set only. **j**, Runtime of all methods on the spleen section.

Human tonsils are organized around B cell follicles embedded within T cell-enriched interfollicular zones, with germinal centers (GCs) forming within follicles to support B cell activation^29^ (**Fig. 3b**, upper left). SpaRank, Cell2location, and RCTD all correctly localized GC cells within annotated GC boundaries and B naive cells in the surrounding follicle regions (**Fig. 3b**). For B plasma cells, all three methods consistently localized predictions to the parenchyma, although SpaRank’s were spatially tighter. Cell2location predicted naïve CD4 T cells broadly across the parenchyma, whereas SpaRank and RCTD yielded more spatially confined distributions. Spatial expression of the canonical marker *CCR7* (**Supp. Fig. S1a**) suggests that this difference may arise from varying sensitivity to marker intensity across methods: Cell2location’s broader predictions overlapped with regions of moderate-to-high CCR7 expression, whereas SpaRank’s predictions overlapped mainly with the highest-expressing spots. This implies that, for this cell type, SpaRank may be more selective for spots with stronger marker signal.

Because distinct tissue compartments typically differ in cellular composition, accurate deconvolution should yield predicted proportions whose clustering recapitulates the underlying anatomical organization. We therefore clustered predicted proportions and evaluated against manual annotations. SpaRank achieved the highest Adjusted Rand Index (ARI) (**Fig. 3c**), with its cluster maps closely recapitulating the annotated compartments—including the connective tissue region that Cell2location failed to resolve (**Supp. Fig. S1b**). Moreover, corresponding anatomical domains across serial sections are expected to share similar cellular compositions, making cross-replicate consistency of predicted proportions a natural measure of algorithmic robustness. We therefore collected three CytAssist Visium tonsil sections with manual anatomical annotations, of which the third was sequenced in a separate batch, introducing pronounced technical variation relative to the first two sections (**Supp. Fig. S1c**). Pairwise Pearson correlation of predicted proportions was then computed within each annotated domain to assess cross-replicate consistency. Tangram achieved the highest mean correlation, with SpaRank closely following and approaching 1 across all domains, demonstrating robust and reproducible inference across technical replicates (**Fig. 3d**).

Within human lymph node, B cell follicles are concentrated in the cortex with T cells enriched in surrounding regions (**Fig. 3e**, upper left). For the CytAssist Visium lymph node section, SpaRank confined GC cell predictions within annotated follicle boundaries, whereas Cell2location and RCTD spread predictions beyond (**Fig. 3e**). *CXCL13*, a key chemokine of the follicular and GC-supporting microenvironment, was spatially restricted to annotated follicles (**Fig. 3f**), corroborating SpaRank’s GC cell localization. Notably, all three methods predicted lower GC proportions than in the tonsil, consistent with the sparse GC activity of normal lymph nodes. For naïve CD4 T cells, SpaRank and RCTD concentrated predictions within the cortex, consistent with CCR7 enrichment in the cortex (**Fig. 3f**), whereas Cell2location extended its predictions into the medullary region. B plasma cells were predicted as medulla-enriched by all methods, though at lower proportions in SpaRank. For the Visium lymph node section, all three methods yielded broadly consistent predictions across cell types (**Fig. 3g**), demonstrating that SpaRank maintains coherent deconvolution performance across sequencing platforms.

Finally, we applied all methods to an Array-seq human spleen section comprising three anatomical regions—white pulp, red pulp, and marginal zone^28^ (**Fig. 3h**). Due to scalability constraints, Cell2location was evaluated on a subsampled set of 7,914 spots, whereas all other methods were evaluated on 51,458 spots (**Methods**). SpaRank predicted germinal center B cells and naive B cells as white pulp-enriched, B plasma cells in the red pulp, and naïve CD4 T cells in the marginal zone (**Fig. 3i**), consistent with the cell-type enrichment reported in the original study^28^. RCTD yielded broadly similar predictions though with lower region specificity, while Cell2location produced less coherent results—including unexpectedly low B naive proportions in the white pulp and B plasma proportions in the red pulp. Notably, SpaRank completed deconvolution in 18 minutes (predominantly pretraining time), compared to 34 minutes for the fastest competing method (SPACEL) and nearly 440 minutes for Cell2location even on the subsampled set (**Fig. 3j**), underscoring the scalability advantage of SpaRank’s pretrain-transfer paradigm.

Overall, pretrained on a comprehensive single-cell atlas derived from human secondary lymphoid organs, SpaRank transfers directly across diverse tissues (tonsil, lymph node, and spleen), anatomical compartments, and sequencing platforms (Visium, CytAssist, Array-seq) without retraining, accurately resolving key cell-type distributions and maintaining cross-replicate consistency.

### SpaRank generalizes across disease states in breast tissue

We next assessed whether SpaRank could generalize across disease states by pretraining on an integrated atlas of normal^30^ and tumor breast tissue^31^ and deploying directly to sections of distinct pathological conditions. This atlas combined normal breast^30^ and breast tumor (spanning ER+, HER2+, TNBC subtypes) ^31^ single-cell datasets, comprising 210,282 cells across 11 major cell types (**Fig. 4a; Methods**). All baseline methods were evaluated under the same single-reference setting.

**Fig. 4.**
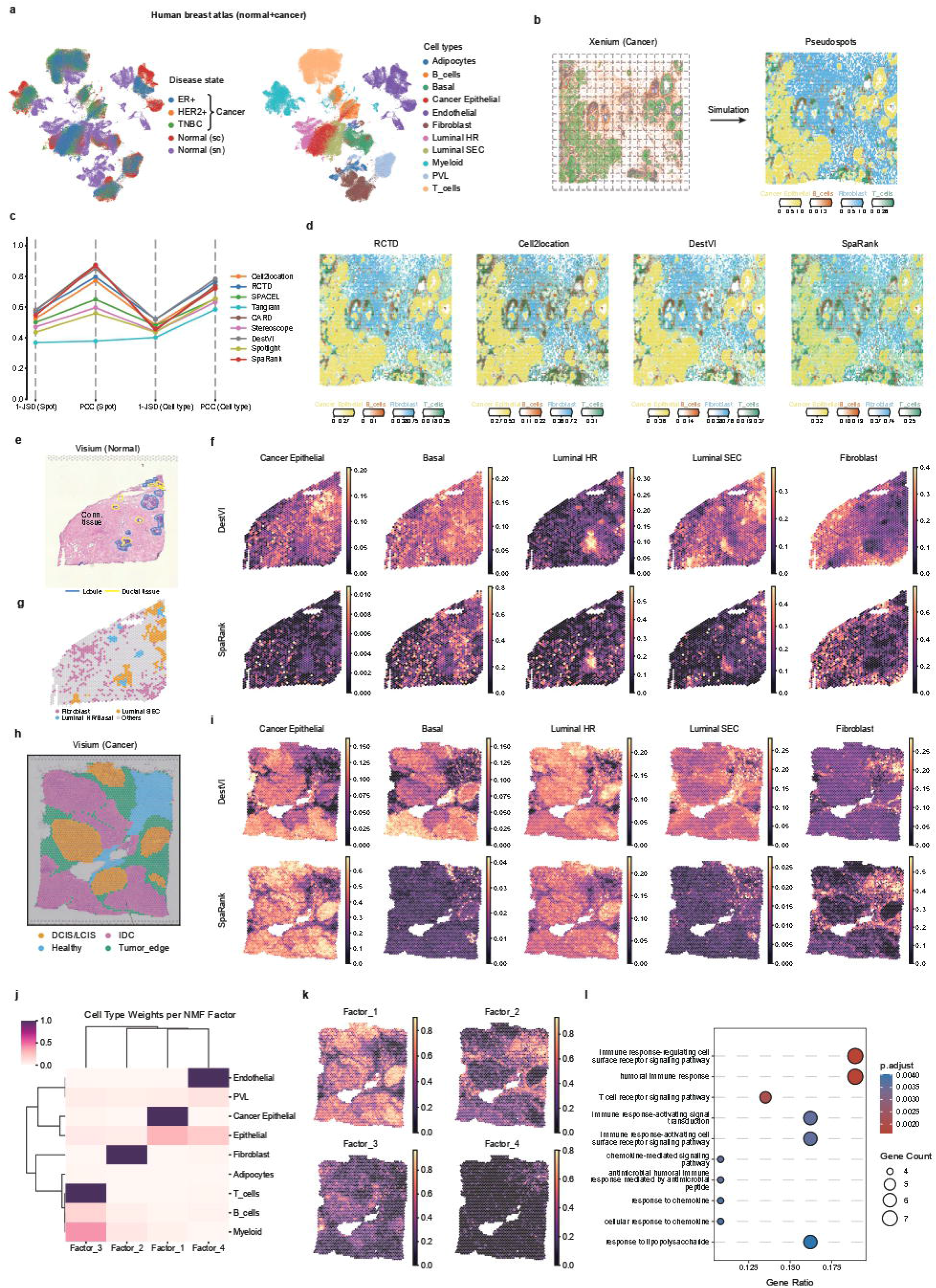
Multi-state transferability across normal and malignant breast tissue. **a**, UMAP visualization of the integrated single-cell reference combining normal and breast cancer tissue, colored by disease state (left) and cell type (right). **b**, Gridding of a Xenium breast cancer section into spot-resolution pseudo-spots (left) and ground-truth abundances of selected cell types (right). **c**, Deconvolution accuracy of all methods on the simulated section, with JSD shown as l − JSD. **d**, Estimated cell-type abundances from four methods on the simulated section. e, H&E image and compartment annotations of the normal Visium section. **f**, Estimated abundances of selected cell types from SpaRank and DestVI on the normal Visium section. **g**, Spot-level cell-type annotations of the normal Visium section from the original study. **h**, Manual anatomical annotations of the Visium breast cancer section shown over the H&E image. i, Estimated abundances of selected cell types from SpaRank and DestVI on the cancer Visium section. **j**, NMF decomposition of SpaRank’s predicted proportions. Heatmap shows estimated factor weights of cell types (rows) across NMF components (columns). **k**, Spatial maps of NMF factor weights. **1**, Gene ontology enrichment analysis of Domain 3, which corresponds to the immune-associated factor.

To establish a quantitative performance baseline under this reference atlas, we gridded an ER+/HER2+ breast cancer Xenium section^32^ into pseudo-spots (**Fig. 4b; Methods**) and benchmarked all methods against ground-truth compositions. SpaRank and CARD achieved the highest spot-level PCC, while on other metrics SpaRank ranked below DestVI and RCTD (**Fig. 4c**). Spatial maps of predicted proportions showed that all methods broadly captured major cell-type distributions, with cancer epithelial dominating large portions of the section and fibroblast as the secondary population (**Fig. 4d**). SpaRank, however, predicted unexpectedly higher T cell proportions in cancer epithelial-enriched regions than RCTD and DestVI.

To demonstrate SpaRank’s transferability across pathological conditions, we analyzed two real Visium sections from distinct contexts—normal breast^30^ and breast cancer tissue^33^. On the normal section (**Fig. 4e**), competing methods predicted substantial cancer epithelial proportions despite the availability of matched normal epithelial populations in the reference atlas, whereas SpaRank assigned very low cancer epithelial proportions (**Fig. 4f**). In addition, SpaRank resolved the spatial distribution of three epithelial subtypes across tissue compartments: Luminal SEC was enriched in lobular regions, while Luminal HR and Basal were concentrated in ductal regions, consistent with spot-level cell-type annotations from the original study^30^ (**Fig. 4g**). All methods agreed on Luminal HR localization, but Cell2location and RCTD additionally predicted substantial Basal proportions in this region (**Supp. Fig. S2a**), whereas the Basal markers KRT14 and KRT17^30^ showed relatively low expression there (**Supp. Fig. S2b**), suggesting over-prediction and supporting SpaRank’s lower Basal estimates. Fibroblast predictions were consistently localized to connective tissue regions across all methods, in agreement with manual annotations (**Fig. 4f,g** and **Supp. Fig. S2a**).

The Visium breast cancer section was of the ER+/HER2+ (Luminal B HER2+) subtype and was manually annotated by pathology assessment of H&E staining^34^ (**Fig. 4h**). All methods predicted cancer epithelial enrichment outside the annotated healthy region, with DestVI yielding the lowest proportions (**Fig. 4i** and **Supp. Fig. S3a**). However, the three epithelial subtypes showed marked inter-method discordance. Specifically, most methods predicted Luminal HR enrichment co-localizing with cancer epithelial predictions in non-healthy regions, consistent with the spatial distribution of *ESR1* and *GATA3*^30^ (**Supp. Fig. S3b**); RCTD, by contrast, predicted low overall Luminal HR proportions concentrated predominantly in DCIS/LCIS regions. This inter-method discordance likely reflects transcriptional similarity between Luminal HR and cancer epithelial reference signatures, both enriched for *ESR1* and *GATA3*, which may confound proportion estimates in tumor regions. For Luminal SEC, SpaRank and RCTD predicted low proportions restricted to healthy regions, supported by uniformly weak *ELF5*^30^ expression across the section and *MFGE8*^35^ expression concentrated in healthy region (**Supp. Fig. S3b**). For Basal cells, DestVI predicted broad enrichment across non-healthy regions, whereas SpaRank, RCTD, and Cell2location confined predictions to annotated tumor edge regions, consistent with *KRT14* and *KRT17* expression restricted to these boundaries (**Supp. Fig. S3b**).

To further characterize the tumor microenvironment captured by SpaRank’s deconvolution, we applied non-negative matrix factorization to the predicted cell-type proportions and extracted four factors representing distinct cellular programs (**Fig. 4j,k**): Factor 1 captured epithelial populations, including both cancer and normal epithelial cells; Factor 2 captured fibroblasts concentrated at annotated tumor edges and in healthy regions; Factor 3 captured T, B, and myeloid cells; and Factor 4 captured endothelial cells enriched in healthy regions. Clustering of the factor weights identified five spatial domains (**Supp. Fig. S3c**). Domain 3 aligned with the immune-associated factor. Differential expression analysis further revealed enrichment of immune-related pathways, including immune response-regulating cell surface receptor signaling and T cell receptor signaling (**Fig. 4l; Methods**), consistent with active anti-tumor immunity. This observation recapitulated findings from a previous study of the same section, which identified a spatially corresponding region of pro-inflammatory immune activity associated with limited tumor growth^34^.

Together, these results demonstrate that a SpaRank model pretrained on an integrated atlas spanning tumor and normal breast tissue can generalize across disease states without retraining. It accurately resolved cell-type distributions in both healthy and tumor sections and enabled biologically meaningful downstream interpretation of the tumor microenvironment.

### SpaRank extends to multimodal spatial deconvolution

The emergence of spatial multi-omics platforms that jointly profile RNA and additional modalities^36,37^ motivates multimodal spatial deconvolution. We therefore used a human thymus single-cell CITE-seq dataset^38^ to construct multimodal benchmark datasets. It comprised 135,566 cells across 38 annotated cell types from 11 batches, with batch effects more pronounced in the RNA modality than in the protein modality (**Fig. 5a**). We constructed 11 benchmark datasets using predefined reference–target batch pairs (**Methods**), rather than a unified reference atlas. This design avoids direct cell-level overlap between reference and target, which would otherwise inflate performance estimates.

**Fig. 5.**
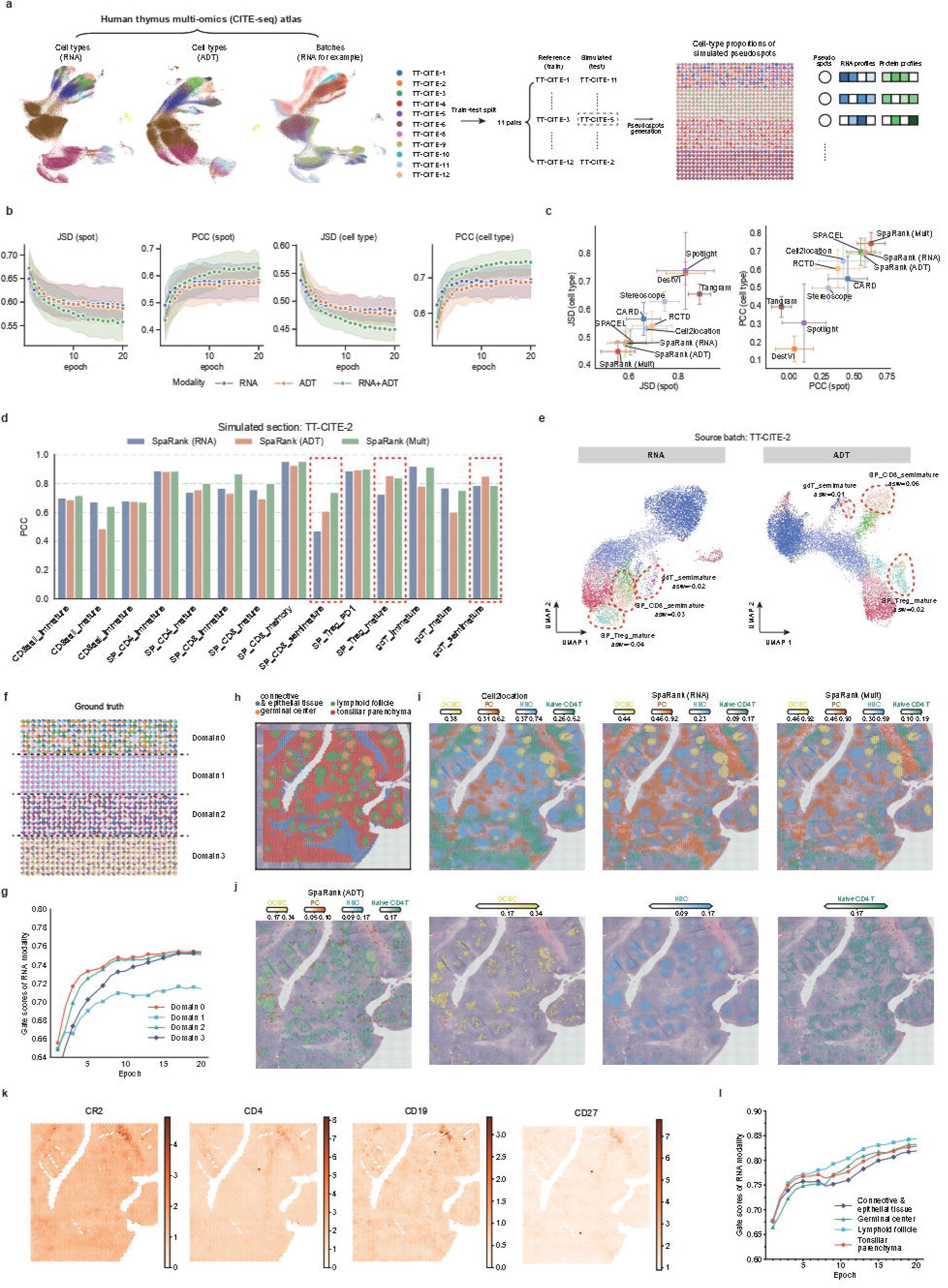
Multimodal spatial deconvolution on simulated thymus and real tonsil data. **a**, UMAP visualization of human thymus CITE-seq data (before integration), colored by cell type in RNA (left), cell type in protein (middle), and batch in RNA (right). Schematic of the benchmark design: reference-target batch pairing and simulation of spatial multi-omics sections from single cells, with ground-truth cell-type compositions shown as spatial pie charts. **b**, Deconvolution accuracy of SpaRank using single-modality data (RNA, ADT) alone and multimodal variants across training epochs. Lines indicate the mean across datasets; shaded regions indicate the interquartile range. **c**, Deconvolution accuracy across **11** simulated datasets, measured by JSD (left) and PCC (right) at the spot level (x-axis) and cell-type level (y-axis). Points indicate the mean; error bars indicate the standard deviation along both axes. **d**, Per-cell-type PCC of SpaRank (RNA), SpaRank (ADT), and SpaRank (Mult) on the TT-CITE-2 simulated section. Red boxes highlight cell types where protein-based predictions exceeded RNA-based predictions. **e**, UMAP visualization of RNA (left) and protein (right) profiles from the TT-CITE-2 source data. Red circles indicate the cell types highlighted in **d;** adjacent values show their silhouette coefficients (ASW). **f**, Ground-truth cell-type abundances of the simulated TT-CITE-2 section, partitioned into four domains by composition. g, Average RNA gate attention scores across the four domains over training epochs on the TT-CITE-2 section. **h**, Manual anatomical annotations of the CytAssist Visium tonsil section. **i**, Estimated cell-type abundances of selected cell types from Cell2location,SpaRank (RNA), and SpaRank (MuJt), shown over the H&E image. **j**, Estimated cell-type abundances from SpaRank (ADT), with GCBC, NBC, and naïve CD4 T cells shown separately for clarity. **k**, Spatial maps of selected protein expression levels. **l**, Average RNA gate attention scores across the four annotated domains over training epochs on the tonsil section.

Training curves across epochs show that SpaRank trained on RNA alone (referred as “SpaRank (RNA)”) and protein alone (referred as “SpaRank (ADT)”) achieved comparable performance across metrics, while joint training on both modalities (referred as “SpaRank (Mult)”) consistently improved performance (**Fig. 5b**). This indicates that the two modalities provide complementary information that SpaRank effectively integrates for improved cell-type deconvolution. Compared with competing methods using RNA alone, SpaRank (RNA) and SpaRank (ADT) performed comparably to SPACEL and ranked among the top-performing approaches (**Fig. 5c**). Integrating both modalities further improved performance, with SpaRank (Mult) achieving the best overall results.

To further dissect modality-specific contributions, we examined predictions on an example simulated section containing 14 cell types (**Fig. 5d**). In terms of cell-type-level PCC, SpaRank (Mult) outperformed both single-modal variants in half of all cell types, while SpaRank (RNA) generally exceeded SpaRank (ADT) across the majority of types, suggesting that RNA provides the dominant signal for distinguishing cell types in this dataset. For those cell types where protein-based predictions surpassed RNA-based ones, such as SP_CD8_semimature, SP_Treg_mature, and gdT_semimature, UMAP visualization of the corresponding single-cell source data revealed that these cell types exhibited sharper cluster boundaries in the protein modality than in RNA (**Fig. 5e**). Also, the silhouette coefficients (ASW) confirmed higher inter-cluster separation for these types in the protein space, demonstrating that SpaRank effectively leverages the discriminative information of each modality across cell types.

To interpret how SpaRank (Mult) weighs each modality, we examined the gate scores assigned to RNA and protein across four spatial domains with distinct compositions (**Fig. 5f and Supp. Fig. S4a**). For each spot, RNA and protein gate scores sum to 1; averaged across spots within each domain, RNA gate scores consistently exceeded protein scores, ranging from 0.7 to 0.8 (**Fig. 5g**), reflecting the dominant role of RNA modality in the model. Domain 1, however, exhibited slightly lower RNA gate scores than the remaining domains (**Fig. 5g** and **Supp. Fig. S4b**), possibly because the protein modality provided a more pronounced inter-cluster separation advantage for its constituent cell types than for those in the other domains (**Supp. Fig. S4c**).

Finally, we evaluated SpaRank on the real CytAssist Visium human tonsil section introduced earlier (**Fig. 5h**), using a human tonsil CITE-seq atlas^39^ comprising 37,119 single cells across 23 major cell types as reference. For comparison, Cell2location was evaluated under the same setting using the RNA modality alone. SpaRank (RNA) and Cell2location correctly localized germinal center B cells (GCBC) within germinal centers, naïve B cells (NBC) in the surrounding follicle, and plasma cells and naïve CD4 T cells in distinct parenchymal regions (**Fig. 5i**). However, SpaRank (ADT) identified NBC but over-predicted GCBC beyond germinal center boundaries and incorrectly distributed naïve CD4 T cells into follicular regions (**Fig. 5j**). This likely arises from the limited cell-type resolution of the ADT panel and the weak spatial specificity of protein measurements captured by the sequencing platform, which reduces its ability to resolve cell-type localization across tonsillar compartments (**Fig. 5k** and **Supp. Fig. S5**). Despite this, SpaRank (Mult) successfully captured the localization of major cell types, consistent with Cell2location and SpaRank (RNA). This was further reflected in gate scores across the four annotated domains, consistently approaching 0.9 for RNA (**Fig. 5l**), indicating that SpaRank adaptively down-weighted the uninformative protein signal.

Together, SpaRank extends naturally to multimodal spatial deconvolution, outperforming single-modality approaches by integrating information across omics layers. Its gated attention mechanism adaptively weights each modality, leveraging informative signals while suppressing noisy ones.

## Discussion

Spatially resolved transcriptomics now enable the mapping of cell types to their native tissue coordinates, revealing how cellular organization and local interactions shape tissue function. Realizing this potential requires computational methods that can accurately and efficiently infer cell-type composition from spatial data across tissues and platforms. Yet existing deconvolution approaches typically require a context-matched single-cell reference and must be retrained for new spatial sections, a paradigm that becomes increasingly impractical as spatial datasets grow in section size, scale, and diversity.

Here, we present SpaRank, a pretrain-transfer framework in which a model trained on a multi-context single-cell atlas can be deployed directly to new spatial sections from the represented contexts without retraining, enabling scalable and generalizable deconvolution across tissues, platforms, and disease states. Quantitative benchmarking on simulated datasets confirmed SpaRank’s accuracy across diverse reference configurations. Furthermore, we evaluated SpaRank in two settings that assess real-world transferability: deploying a model pretrained on a multi-organ lymphoid atlas to sections spanning distinct tissue origins and sequencing platforms, and deploying a model pretrained on an integrated breast reference spanning tumor and normal tissue to sections representing distinct pathological states. In both settings, SpaRank accurately resolved key cell-type distributions, indicating robustness to real-world variation.

Since SpaRank imposes no distributional assumptions on input data, it naturally extends to multimodal spatial deconvolution through a gated multimodal fusion mechanism. On simulated sections, SpaRank trained jointly on RNA and protein outperformed its single-modality variants and competing methods; on a real CytAssist tonsil section where protein measurements lacked spatial specificity, SpaRank maintained accurate localization by adaptively suppressing the noisy protein modality. What’s more, SpaRank’s framework confers substantial scalability advantages: deconvolution of a large Array-seq spleen section was completed within 18 minutes, compared to nearly 440 minutes for Cell2location even on a subsampled set.

Future development of SpaRank will focus on scaling and architectural refinement. Pretraining on more comprehensive single-cell atlases will establish universal deconvolution models capable of resolving highly complex biological contexts. Concurrently, integrating explicit spatial priors into the architecture will leverage tissue topology to regularize predictions and mitigate the localized noise inherent to independent spot-wise predictions.

## Methods

### Simulation of ST data from single-cell data

To generate training data for SpaRank, a total of *N* pseudo-spots are simulated by sampling cells from the single-cell reference, with each spot containing cells from only one biological context, such as a tissue, organ, or disease state. The *N* spots are allocated across contexts in proportion to their respective cell counts, preserving the compositional diversity of the reference. For each simulated spot, the number of constituent cells *N*_*c*_ and the number of constituent cell types *N*_*t*_ are sampled independently from either a normal distribution 𝒩(*μ*_*c*_, *σ*_*c*_) and 𝒩(*μ*_*t*_, *σ*_*t*_), or a log-normal distribution LogN (*μ*_*c*_, *σ*_*c*_) and LogN (*μ*_*t*_, *σ*_*t*_), the latter better capturing the overdispersed cellular densities observed in complex tissue microenvironments.

Once *N*_*c*_ and *N*_*t*_ are determined, *N*_*t*_ cell types are selected from the reference according to a cell-type sampling distribution *P*_*t*_. Following SPACEL^12^, three sampling regimes are applied in equal proportion to generate spots with diverse cell-type compositions, each governed by a uniform random variable *r*_*c*_:

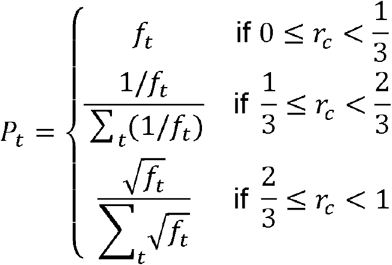

where *f*_*t*_ is the observed frequency of cell type *t* in the reference. When 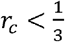, cell types are sampled proportionally to their observed frequencies, preserving the natural compositional imbalance of the reference. When 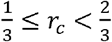, all cell types are sampled with equal probability, encouraging uniform representation. When 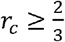, a square-root transformation down-weights abundant cell types while up-weighting rare ones, balancing the two. The expression profile of each simulated spot is then obtained by summing the modality expression profiles of all *N*_*c*_ sampled cells. The ground-truth cell-type proportion label for each simulated spot is computed by counting the number of sampled cells belonging to each cell type and normalizing by *N*_*c*_, yielding a compositional vector **y** ∈ ℝ^K^ where *K* is the number of total cell types in the reference data and ∑_*k*_ *y*_*k*_ = 1.

### Feature selection and input tokenization

Single-cell RNA-seq reference datasets typically profile tens of thousands of genes, the majority of which are uninformative for cell-type discrimination and would impose unnecessary computational burden if used directly as model input. To identify a compact and discriminative gene set, differentially expressed genes (DEGs) are selected from the single-cell reference using a batch-wise Wilcoxon rank-sum test, retaining genes with log_2_ fold-change ≥ 0.5 and adjusted *p*-value <0.01 in at least one batch. Alternatively, the input gene set may be specified as a priori. For example, in our experiments, spot-resolution sections simulated from Xenium and MERFISH data inherited the corresponding platform-defined gene panels, which were used directly as the input gene set. In either case, the final gene set was obtained by intersecting with the genes present in the single-cell reference, thereby ensuring compatibility between reference and query.

The selected features define a vocabulary 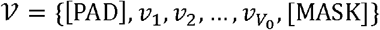 of size *V* = *V*_0_ + 2, with special tokens for padding [PAD] and masking [MASK]. Each feature is assigned a unique integer identifier within this vocabulary. For each spot, expression values are ranked in descending order, and the top *K*_*g*_ features are extracted as an ordered token sequence 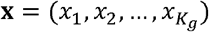, where each *x*_*i*_ ∈ *V*. Positions are padded with [PAD] when fewer than *K*_*g*_ features are expressed in a given spot.

In the multimodal setting, feature selection and vocabulary construction are performed independently per modality, yielding modality-specific vocabularies *V*^*m*^, token sequences **x**^*m*^, and sequence lengths 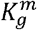. For protein, all measured features are retained without filtration.

### Model architecture

We first describe SpaRank in the single-modality setting, followed by its multimodal extension. Given the input token sequence 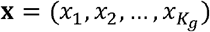 defined above, each token is mapped to a dense vector via a learnable embedding table *E* ∈ ℝ ^| 𝒱| ×*d*^, where *d* is the embedding dimension. A learnable [CLS] token **e**_cls_ ∈ ℝ^*d*^ is prepended, and learnable positional embeddings 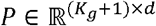 are added to encode rank order, yielding the input matrix 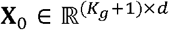. This matrix is passed through *L* Transformer encoder layers, each comprising multi-head self-attention with *H* heads and a feed-forward network of dimension *d*_ff_, with layer normalization^40^ and dropout^41^ applied throughout. The output at the [CLS] position serves as the spot-level embedding **h** ∈ ℝ^*d*^, and the outputs at the remaining token positions are denoted **t**_*i*_ ∈ ℝ^*d*^ for *i* = 1, … *K*_*g*_.

To condition predictions on the biological context of the reference, such as tissue type or disease state, a learnable context embedding 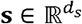 is concatenated to **h** after the encoder, yielding a fused Representation 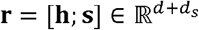. This fused representation is passed to two heads: (1) a prediction head 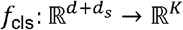, implemented as a two-layer multi-layer perceptron (MLP) with Gaussian Error Linear Unit (GELU) activation^42^, that outputs cell-type proportion logits over *K* cell types; and (2) a projection head 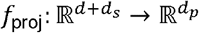 that maps **r** to a lower-dimensional space for contrastive learning where *d*_*p*_ = 64. A third head *f*_rec_: ℝ^*d*^ → ℝ^*V*^ operates on the per-token encoder outputs and predicts the identity of masked tokens during pretraining.

For multimodal deconvolution, each modality *m* ∈ {1, …, *M*} is encoded by an independent Transformer encoder following the same architecture described above, with a dedicated reconstruction head 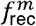. The modality-specific CLS embeddings **h**^*m*^ ∈ ℝ^*d*^ from each encoder are integrated via a gated fusion mechanism. A gate vector for each modality is computed as:

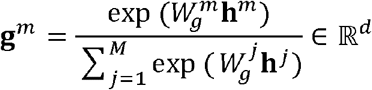

where 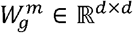 are learnable projection matrices and the formulation ensures 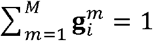 for every dimension *i*. The fused multi-modal embedding is then obtained as 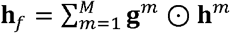, where ⊙ denotes element-wise multiplication. This fused embedding is then concatenated with the context embedding and passed to the shared prediction and projection heads *f*_cls_ and *f*_proj_.

### Pretraining

A central challenge in training SpaRank is the risk that the model overfits to top-ranked tokens, which represent features with high expression and strong cell-type specificity. These provide a brittle learning signal, as such features may be undetected or captured at much lower levels in the target spatial data. To mitigate this, SpaRank applies stochastic token masking as a core regularization strategy throughout training.

For each spot, given an input token sequence **x** per modality, three independent masking operations are applied: two random token dropout operations that each set a fraction *p*_drop_ of non-padding tokens to the padding identifier, producing views **x**^(*a*)^ and **x**^(*b*)^, and one random masking operation that replaces a fraction *p*_mask_ of non-padding tokens with a [MASK] identifier to produce 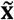, with the masked positions forming the set ℳ. In the multimodal setting, masking is applied independently to each modality with modality-specific rates, 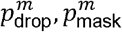.

#### Contrastive loss

The two views **x**^(*a*)^ and **x**^(*b*)^ are each passed through the encoder to obtain spot-level embeddings **h**^(*a*)^ and **h**^(*b*)^. In the single-modality setting, these are concatenated with the context embedding **s** to form **r**^(*a*)^ = [**h**^(*a*)^; **s**] and **r**^(*b*)^ = [**h**^(*b*)^; **s**]. In the multimodal setting, the modality-specific embeddings are first integrated via gated fusion to obtain 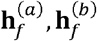, and **r**^(*a*)^, **r**^(*b*)^. The projection head maps each to **z**^(*a*)^ = *f*_proj_ (**r**^(*a*)^) and **z**^(*b*)^ = *f*_proj_ (**r**^(*b*)^), and the contrastive learning loss is computed as^19^ :

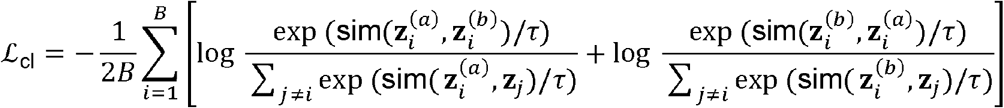

where sim (·, ·) denotes cosine similarity, *τ* is a temperature parameter, and *B* is the batch size. This loss discourages the encoder from relying on a fixed set of top-ranked tokens.

#### Reconstruction loss

The masked sequence 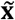 is passed through the encoder, and the per-token outputs at masked positions are fed to the reconstruction head *f*_rec_ to predict the original token identities:

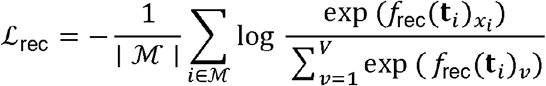

Where **t**_*i*_ is the encoder output at position *i*; *f*_rec_ (**t**_i_) ∈ ℝ^*V*^ are the predicted logits and *x*_i_ indexes the ground-truth token. This encourages the encoder to capture global co-expression dependencies across the full sequence. In the multimodal setting, reconstruction is performed independently per modality through dedicated heads 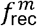, and the per-modality losses are averaged: 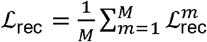.

#### Composition prediction loss

The representation **r**^(*a*)^ from the first augmented view is passed to the prediction head to obtain cell-type proportion logits **ŷ** = *f*_cls_ (**r**^(*a*)^). A cross-entropy loss is computed against the ground-truth proportions **y**:

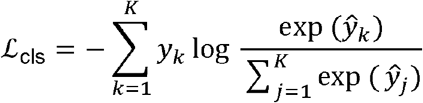

The total training objective is:

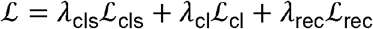

where *λ*_cls_, *λ*_cl_, and *λ*_rec_are scalar weights.

### Inference

At inference, the vocabulary set is applied to filter and rank expressed features in each spot of the target spatial section, following the same tokenization procedure described above. A state identifier is assigned to each spot according to the biological context of the target section. The token sequence and state identifier are passed through the Transformer encoder and head networks to produce cell-type proportion estimates directly.

### Implementation Details

For simulation of datasets used for pretraining, *N*_*c*_ and *N*_*t*_ were sampled from either normal or log-normal distributions depending on the dataset. The mouse isocortex dataset used log-normal distributions with *μ*_*c*_ = 15, *σ*_*c*_ = 10, and *N* = 1,000,000 pseudo-spots. The human thymus dataset used normal distributions with *μ*_*c*_ = 10, *σ*_*c*_ = 5, and *N* = 500,000 pseudo-spots. Remaining datasets—human lymphoid organs, human breast, and multimodal tonsil experiments—used normal distributions with *μ*_*c*_ = 10, *σ*_*c*_ = 5, and pseudo-spots, except human breast which used log-normal distributions with the same parameters. All sampling was clipped to [1,50] across all datasets.

SpaRank was implemented in PyTorch^43^ and optimized using AdamW^44^ with learning rate 1 ×10^−4^, batch size 256, and gradient clipping with maximum norm 1.0. The loss weighting coefficient for the composition prediction loss was fixed at *λ*_cls_ = 1.0 across all experiments. The contrastive and reconstruction losses were applied in all experiments except the mouse isocortex dataset, where *λ*_cl_ = *λ*_rec_ = 0 was used; for all other datasets, *λ*_cl_ = 0.5 and *λ*_rec_ 0.5 with temperature *τ* = 0.1. Token dropout rate *p*_drop_ = 0.3 and masking rate *p*_mask_ = 0.3 were set across all experiments and modalities. The input token sequence length for RNA was set to *K*_*g*_ = 500 for all experiments. For the multimodal deconvolution experiments, 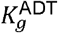 was set to the number of measured proteins.

For the mouse isocortex and human thymus simulation datasets, a larger Transformer encoder was used: *L* = 2, *H* = 4, *d* = 128 and *d*_ff_ = 256. For all real datasets, *L* = 1, *H* = 1, *d* = 64 and *d*_ff_ = 256 were used to avoid overfitting. The state embedding dimension was set to *d*_*s*_ = *d*/8 in all experiments. All parameters were initialized using Xavier uniform initialization^45^.

### Single-cell reference atlas used in analysis

#### Mouse isocortex

Single-cell RNA-seq data were obtained from the Allen Institute whole-brain atlas^21^, profiled using 10x Chromium v2 and v3 chemistries; only data generated with 10x v3 were used here. We focused on the isocortex region, which encompasses 24 cell-type classes with well-defined cytoarchitectural organization. To reduce dataset size, cell types exceeding 100 cells were downsampled to 10% of their original count, yielding a reference of 46,575 cells across 32,285 genes. To simulate perturbations to reference expression profiles, we applied a gene-specific nonlinear transformation and stochastic dropout to the original count matrix. Specifically, for each gene *g*, a power factor *p*_*g*_ was independently sampled from a uniform distribution, *p*_*g*_ ~U(max (0.5, 1 −0.2*λ*),1+0.2*λ*), to introduce gene-wise non-linear rescaling of expression magnitudes, where *λ* denotes the perturbation intensity. An efficiency factor *e*_*g*_ was independently sampled from a log-normal distribution, *e*_*g*_ ~LogNormal(0, 0.2, *λ*), to mimic gene-specific amplification biases across platforms. The perturbed count for each non-zero entry was computed as: 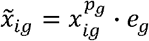. To further simulate transcript dropout, each perturbed value was stochastically set to zero using a logistic retention probability: 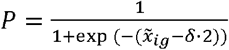 where *δ* is the sparsity loss coefficient, such that low-abundance transcripts are preferentially dropped. Final values were rounded to the nearest integer to preserve count values. Five perturbation levels were defined spanning from raw (*λ* = 0, *δ* = − ∞) to level 4 (*λ* = 3.5, *δ* = 0.6) **(Supp. Fig. S6**).

#### Human lymphoid organs

The single-cell reference atlas was obtained from Kleshchevnikov et al.^6^, comprising scRNA-seq datasets from human lymph nodes, spleen, and tonsils integrated across three independent studies^23-25^. Batch effects were corrected using BBKNN^46^, yielding 73,260 cells across 34 annotated cell populations.

#### Human breast

The single-cell reference was constructed by integrating two publicly available atlases: an atlas of human breast cancers spanning ER+, HER2+, and TNBC subtypes^31^, and an atlas of normal adult human breast tissue profiled by both scRNA-seq and snRNA-seq^30^. For the normal atlas, scRNA-seq and snRNA-seq data were independently downsampled to at most 6,000 cells per cell type, yielding 57,720 and 52,498 cells respectively. The two atlases were merged and preprocessed by selecting highly variable genes, log-normalization, and PCA, followed by Harmony^47^ integration across all batches. After harmonizing cell type annotations, the final reference comprised 11 major cell types across 210,282 cells.

#### Human thymus

A human thymus single-cell CITE-seq dataset^38^ was used to construct multimodal benchmark datasets. It comprised 135,566 cells across 38 annotated cell types from 11 batches. To construct benchmark pairs, each batch was assigned to its most compositionally similar counterpart based on pairwise cell-type composition similarity, yielding 11 pairs with comparable cell-type compositions. Within each pair, one batch served as the single-cell reference and the other was used to simulate the spatial section.

### Simulated and real spatial sections used in analysis

#### Mouse isocortex

We collected a single-cell resolution MERFISH atlas of the adult mouse brain^20^ comprising approximately 10 million cells profiled across 1,100 genes, with cell-type labels transferred from the Allen Institute whole-brain scRNA-seq atlas described above. We focused on the isocortex region, retaining 27 sections with at least 5,000 isocortex cells. To simulate spot-resolution spatial data, a uniform spatial grid was applied to each section using a window size of 0.12 (in scaled coordinates) to aggregate cells into pseudo-spots with ground-truth cell-type proportions. The window size was chosen to yield spots at a density comparable to that of 10x Visium.

#### Human tonsil

Three CytAssist Visium human tonsil replicate sections were collected at 55 *μ*m resolution, each with manual anatomical annotations and paired RNA and protein measurements^48^. For the cross-organ transferability analysis, only the RNA modality was used. For the multimodal deconvolution analysis, both RNA and protein measurements from the first replicate were used. As the protein panel nomenclature differed between the single-cell CITE-seq reference and the CytAssist section, protein names were harmonized across the two datasets, yielding 25 shared proteins.

#### Human lymph node

A CytAssist Visium human lymph node section^48^ and a publicly available standard Visium lymph node section were collected^27^, both at 55 *μ*m resolution. The CytAssist section carries manual anatomical annotations and paired RNA and protein measurements; only the RNA modality was used in the cross-organ transferability analysis.

#### Human spleen

The Array-seq spleen section^28^ originally comprised 750,640 spots. As most methods required prohibitive runtimes on the full section, we cropped a central region of 51,458 spots for benchmarking. Even at 51,458 spots, Cell2location required hundreds of hours and suffered numerical instability, so we further cropped to a central subset of 7,914 spots.

#### Human breast

We obtained an ER+/HER2+ Xenium breast cancer section comprising 158,030 single cells profiled across a 313-gene panel^32^. Cell-type labels were harmonized to match the annotations of the breast cancer single-cell reference described above, with ambiguous hybrid cells removed, yielding 8 major cell types^49^. To mimic Visium spot resolution, cells were aggregated into pseudo-spots by binning within a 55 µm square grid, summing gene expression profiles of all cells within each bin while preserving ground-truth cell-type compositions. The normal breast Visium section was obtained from a normal adult human breast atlas, which provides manual annotations of epithelial compartments including connective tissue, ductal, and lobular regions. The tumor breast Visium section was obtained from the publicly available 10x Genomics dataset repository^33^. Xu et al.^34^ annotated tumor and healthy domains on this section based on H&E histological assessment.

#### Human thymus

Following the simulation framework of Sarah et al.^50^, pseudo-spots with paired RNA and ADT (protein) expression profiles were generated by randomly sampling and aggregating cells from a multimodal single-cell reference. For each batch, cell types with fewer than 15 cells were excluded prior to simulation. A grid of 1,000 pseudo-spots was arranged on a 2D coordinate system and partitioned into 4 spatial stripe regions, each assigned a distinct set of 3–5 cell types drawn without replacement under balanced sampling. This ensured equal selection probability across all cell types, regardless of their abundance in the reference. The number of cells per spot was drawn from a Conway–Maxwell–Poisson distribution with region-specific means (5, 10, or 15 cells) and a fixed dispersion parameter, 20. For each pseudo-spot, the sampled cells’ RNA and ADT count profiles were summed to produce the aggregated expression profile, and the ground-truth cell type composition was recorded.

### Differential expression analysis and GO enrichment analysis

We used the Wilcoxon test, as implemented in the rank genes groups function of the SCANPY^51^ package (version 1.9.5), to identify differentially expressed genes of spatial clusters. We performed the GO enrichment analysis on cluster-specific differentially expressed genes using the R package clusterProfiler^52^ (version 4.6.2)

### Compared methods

#### Cell2location

Cell2location models cell-type composition in spatial transcriptomics data through a Bayesian negative binomial regression framework^6^. We used version 0.1.4 and followed the guidelines from https://github.com/BayraktarLab/cell2location, with default parameters except for n_cells_per_location, which was set to 30 for all Visium sections and for simulated sections derived from the HER2+ Xenium breast cancer sample, and to 10 for simulated sections derived from the human thymus and mouse isocortex datasets. Cell-type abundances were estimated from the means_cell_abundance_w_sf posterior.

#### CARD

CARD is a conditional autoregressive deconvolution method that jointly models cell-type-specific expression and spatial correlation in cell-type compositions across tissue locations^9^. We used version 1.1 and followed the guidelines from https://yma-lab.github.io/CARD/, with default parameters set for all datasets.

#### DestVI

DestVI deconvolves spatial transcriptomics data through a two-stage variational inference framework, first learning cell-type-specific latent representations from a scRNA-seq reference and then inferring spatial cell-type proportions^8^. We used scvi-tools^53^ v1.0.3 and followed guidelines from https://docs.scvi-tools.org/en/stable/tutorials/notebooks/spatial/DestVI_tutorial.html, with max_epochs set to 300 for the scLVM and 2000 for the stLVM, with all other parameters set to default.

#### RCTD

RCTD infers cell-type proportions using maximum-likelihood estimation under a Poisson count model with a log-normal prior^10^. We used the spacexr package v2.2.1 and followed the guidelines from https://raw.githack.com/dmcable/spacexr/master/vignettes/visium_full_regions.html, with doublet_mode set to ‘full’ and filtered cell types with fewer than 25 cells from the reference. Predicted weights were normalized using the built-in normalize_weights function, with all other parameters set to default. For the human breast dataset, whose reference atlas comprises over 200,000 cells, we used an optimized Python implementation (https://github.com/p-gueguen/rctd-py) with identical parameter settings.

#### SPOTlight

SPOTlight deconvolves spatial transcriptomics data using seeded non-negative matrix factorization (NMF) initialized with cell-type marker genes, followed by non-negative least squares (NNLS) regression^11^. We used version 1.6.7 and followed the guidelines from https://marcelosua.github.io/SPOTlight/articles/SPOTlight_kidney.html. We set the marker gene AUC threshold to 0.65, the minimum cell-type proportion to 0.01, and the number of cells sampled per cell type to 100, with all other parameters set to default.

#### Stereoscope

Stereoscope is a probabilistic deconvolution method that trains two sequential latent variable models: a single-cell reference model that learns cell-type-specific expression parameters from scRNA-seq data, and a spatial model that incorporates the learned reference parameters to infer cell-type proportions in spatial transcriptomics data^7^. We used scvi-tools v1.0.3 and followed guidelines from https://docs.scvi-tools.org/en/1.3.3/tutorials/notebooks/spatial/stereoscope_heart_LV_tutorial.html. The reference and spatial models were trained for 100 and 2,000 epochs respectively, with all other parameters set to default.

#### SPACEL

SPACEL deconvolves cell-type composition for each spatial spot using a multilayer perceptron combined with a probabilistic model^12^. We used version 1.1.8 and followed the guidelines from https://github.com/QuKunLab/SPACEL/blob/main/docs/tutorials/Visium_human_DLPFC_Spoint.ipynb. We used the Spoint module with DEGs identified by t-test, number of simulated spots st to 100,000, and trained for 5,000 steps with batch size 1,024, with all other parameters set to default.

#### Tangram

Tangram maps single-cell gene expression to spatial locations through non-convex optimization, producing a probability matrix over cell types for each spatial spot^13^. We used version v1.0.4 with default parameters and ran Tangram in ‘cluster’ mode, following the guidelines at https://tangram-sc.readthedocs.io/en/latest/tutorial_link.html.

## Data availability

All datasets used in this study are publicly available. The whole mouse brain single-cell reference atlas is available at https://brain-map.org/bkp/explore/abc-atlas; instructions for accessing the processed scRNA-seq data are provided at https://github.com/AllenInstitute/abc_atlas_access/blob/main/descriptions/WMB-10X.md. The corresponding whole mouse brain MERFISH data are accessible via the CELLxGENE database (https://cellxgene.cziscience.com/collections/0cca8620-8dee-45d0-aef5-23f032a5cf09). The Allen Mouse Brain Common Coordinate Framework (CCFv3) annotations are available at https://alleninstitute.github.io/abc_atlas_access/descriptions/Allen-CCF-2020.html. The curated human lymphoid organ reference atlas and the human lymph node Visium section are accessible through the Cell2location tutorial (https://cell2location.readthedocs.io/en/latest/notebooks/cell2location_tutorial.html). CytAssist Visium sections of lymph node and tonsil are available at https://zenodo.org/records/18946723. The human normal breast reference atlas and paired Visium section are available via CELLxGENE (https://cellxgene.cziscience.com/collections/4195ab4c-20bd-4cd3-8b3d-65601277e731); the breast cancer single-cell reference atlas is deposited in the Gene Expression Omnibus (GEO) under accession GSE176078. The human breast cancer Xenium section is available from GEO under accession GSE243280, with cell-type annotations provided by 10x Genomics (https://www.10xgenomics.com/products/xenium-in-situ/preview-dataset-human-breast). The human breast cancer Visium section is available from 10x Genomics (https://www.10xgenomics.com/datasets/human-breast-cancer-block-a-section-1-1-standard-1-1-0), and the corresponding pathological annotations are available at https://github.com/JinmiaoChenLab/SEDR_analyses/. The human thymus dataset is accessible via CELLxGENE (https://cellxgene.cziscience.com/collections/fc19ae6c-d7c1-4dce-b703-62c5d52061b4). All processed datasets are publicly available at Zenodo (https://zenodo.org/records/20091867).

## Code availability

The SpaRank open-source Python package is available at GitHub (https://github.com/XiHuYan/SpaRank).

## Authors’ contributions

W.L, M.L. and J.C. initiated the project and provided funding support. X.Y. and R.Z. developed the method and designed the experiments. X.Y. and W.L performed the data analysis and wrote the manuscript. All authors reviewed and approved the final manuscript.

## Acknowledgements

W.L is supported by the Natural Science Foundation of Guangxi (2024GXNSFFA010006); the National Natural Science Foundation of China (U24A20256 and 62472108). M.L. is supported in part by the National Natural Science Foundation of China under Grant (No.62225209 to M.L.). J.C. is supported by Ministry of Education Academic Research Fund Tier 2 (CELL2VIRUS: AI-Driven Mapping of Virus-Host Interactions at Single-Cell and Spatial Resolution, MOE-T2EP30125-0005); NMRC Open Fund Large Collaborative Grant (Singapore 1YMPHoma translatiONal studY (SYMPHONY) 2.0, MOH-001575-00).

---

**Supp. Fig. S1.**
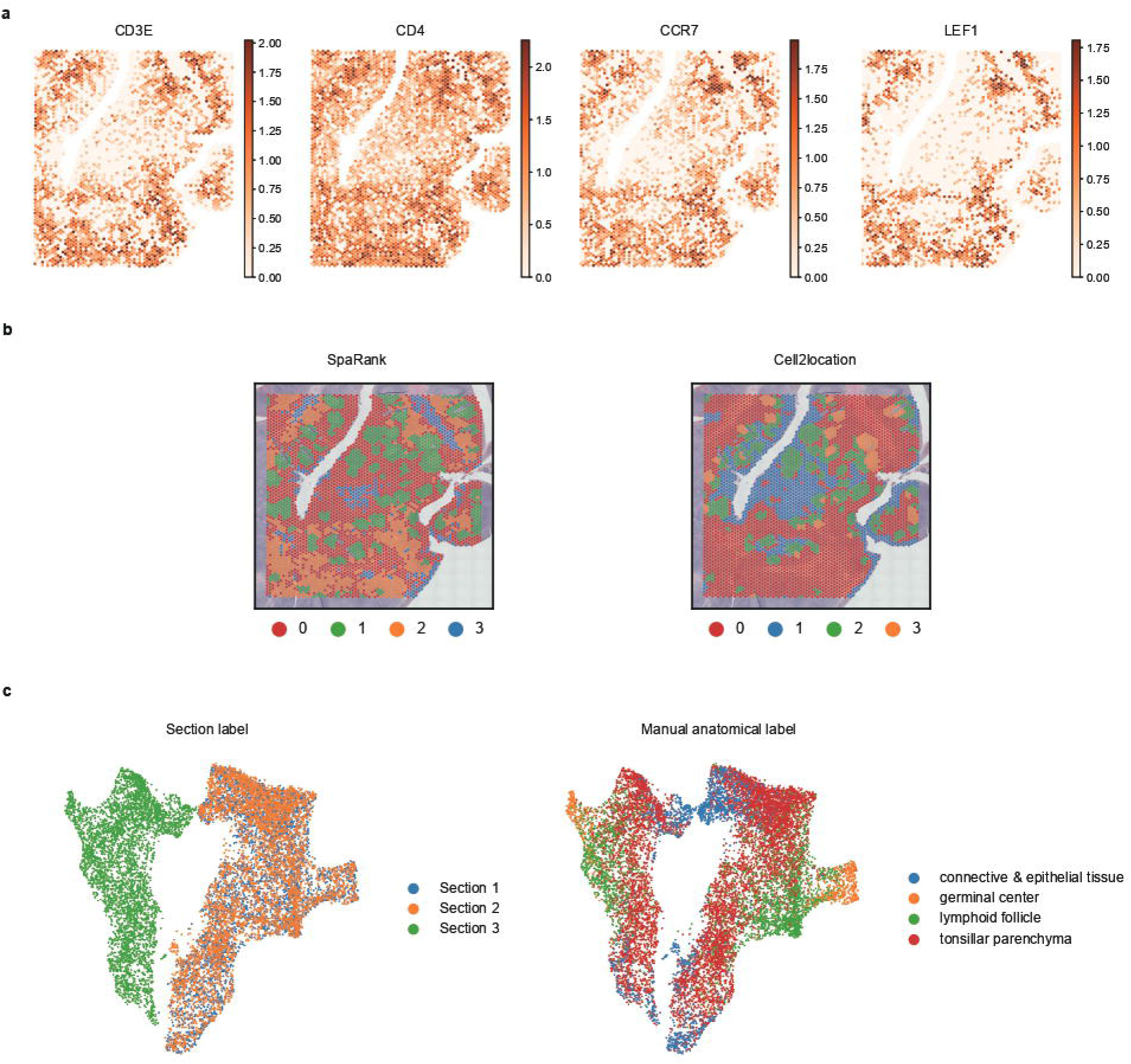
Additional results for the human tonsil section in the multi-organ transferability analysis. **a**, Spatial RNA expression maps of naïve CD4T cell marker genes *CD3D,CD4, CCR7*, and *LEF1*. **b**, Louvain clustering of predicted proportions from SpaRank and Cel12location at a resolution of four clusters. **c**, UMAP visualization of RNA expression across three tonsil replicates, colored by batch (left) and anatomical domain annotation (right).

**Supp. Fig. S2.**
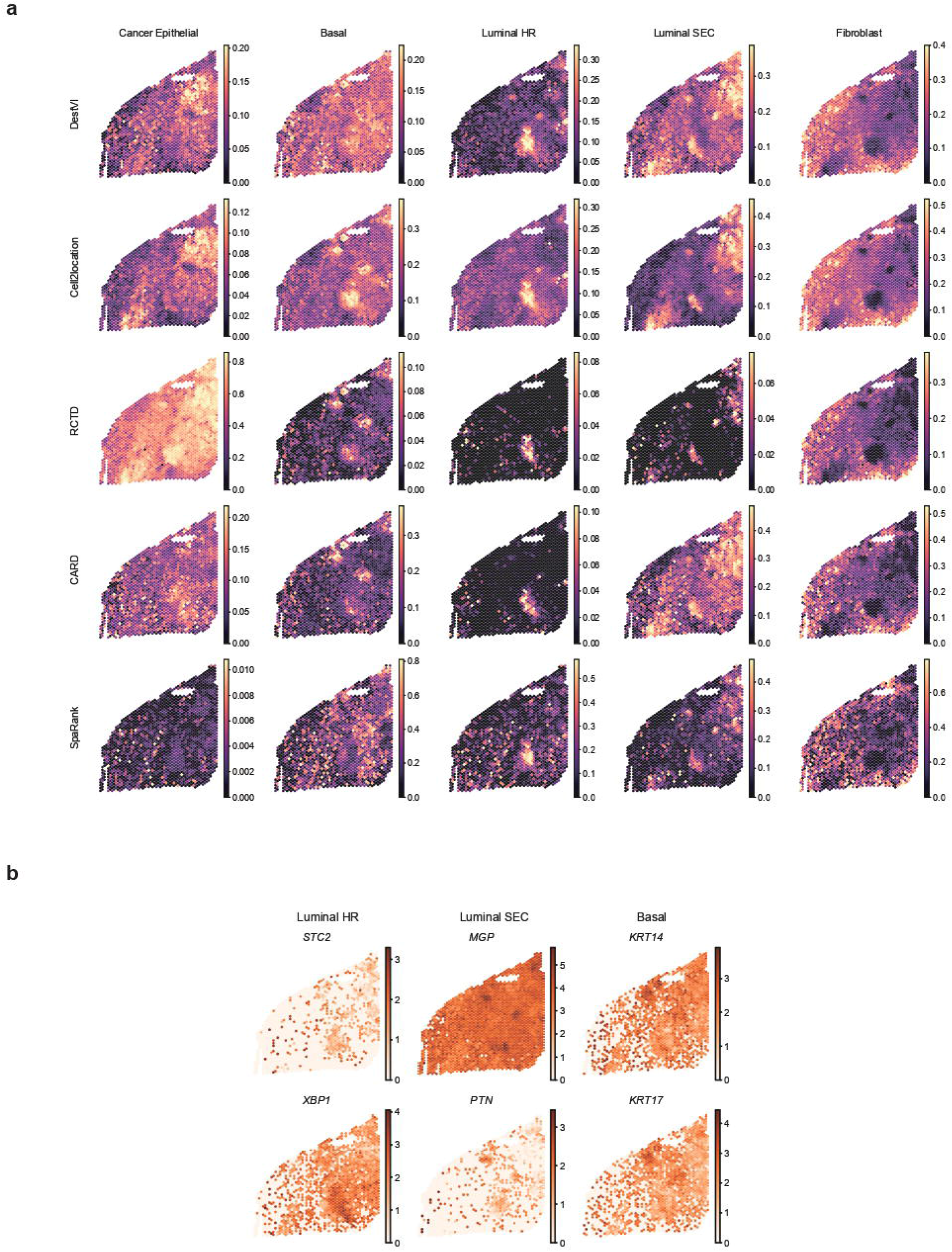
Additional results for the normal Visium breast section in the multi-state transferability analysis. **a**, Estimated cell-type abundances of selected cell types from DestVl, Cell21ocation, RCTD, CARD, and SpaRank. **b**, Spatial RNA expression maps of marker genes for three epithelial subtypes: Luminal HR, Luminal SEC, and Basal.

**Supp. Fig. S3.**
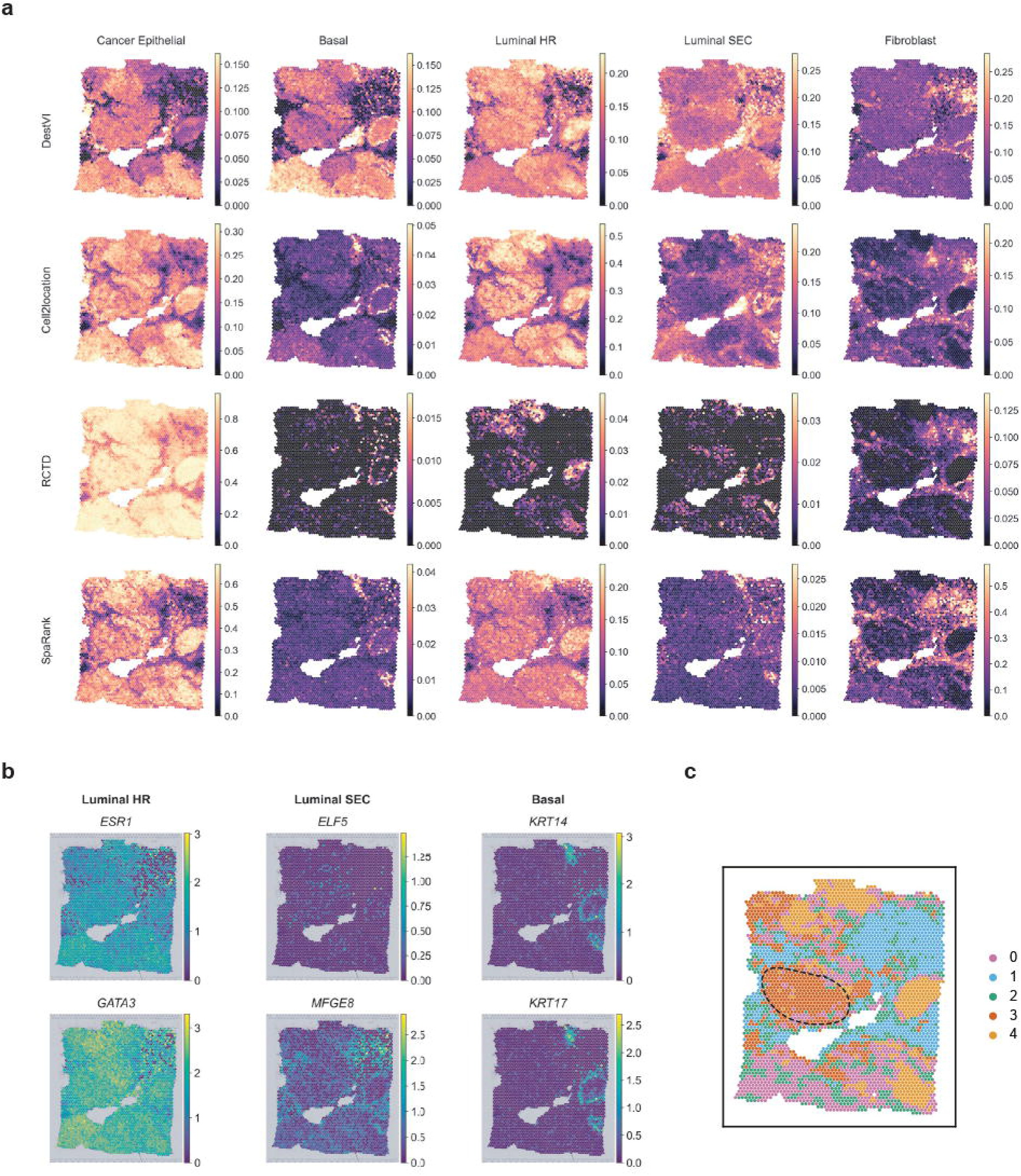
Additional results for the cancer Visium breast section in the multi-state transferability analysis. **a**, Estimated cell-type abundances of selected cell types from DestVI, Cell2location, RCTD, and SpaRank. **b**, Spatial RNA expression maps of marker genes for three epithelial subtypes: Luminal HR, Luminal SEC, and Basal. c, Louvain clustering of NMF factor weights derived from SpaRank’s predicted proportions.

**Supp. Fig. S4.**
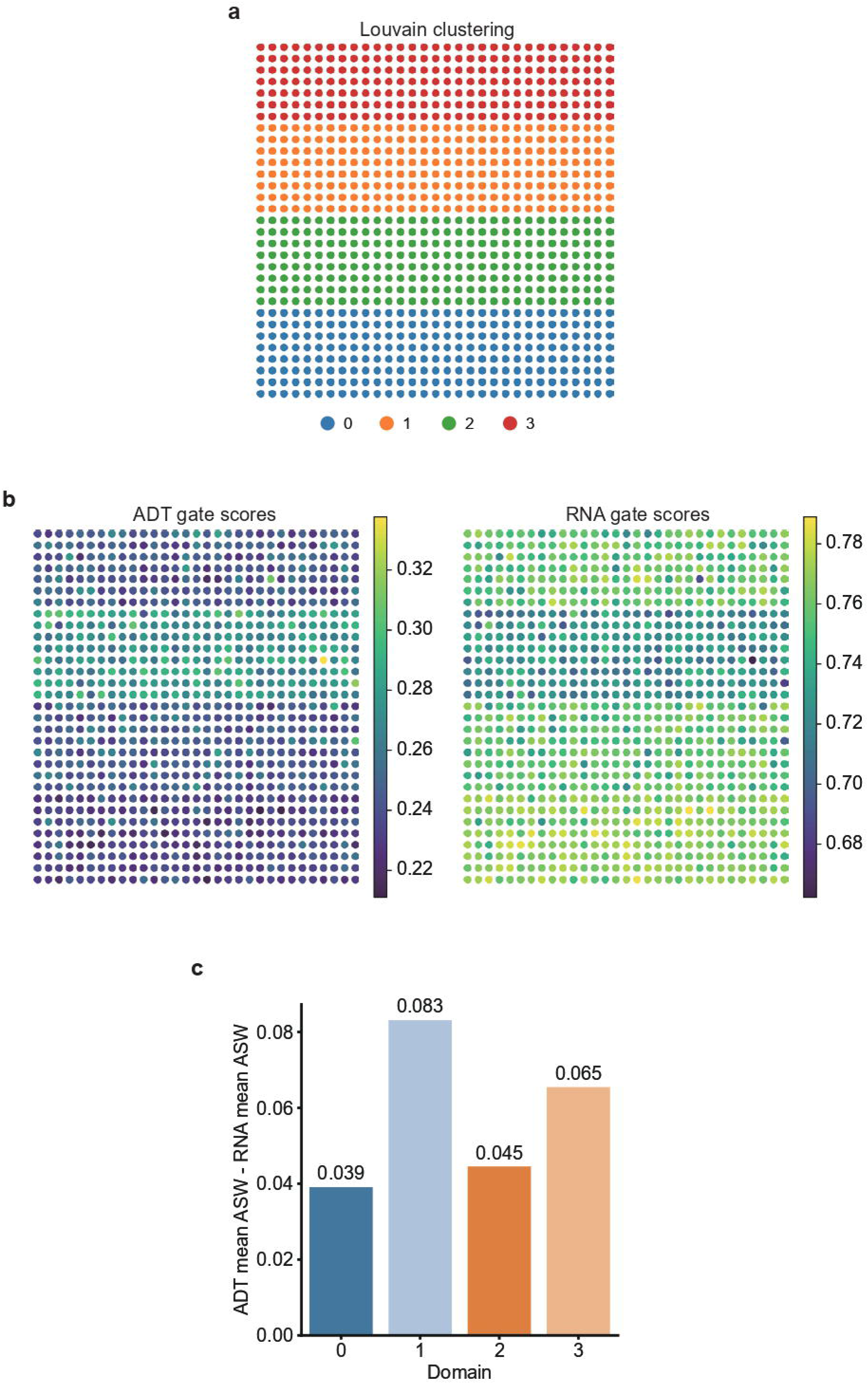
Gate attention analysis on the TT-CITE-2 simulated section. **a**, Louvain clustering of ground-truth cell-type abundances identifying four spatial domains. **b**, Spatial maps of spot-level gate attention scores for the RNA and protein modalities. **c**, Difference in silhouette coefficient (ASW) between protein and RNA modalities, averaged across cell types within each spatial domain.

**Supp. Fig. S5.**
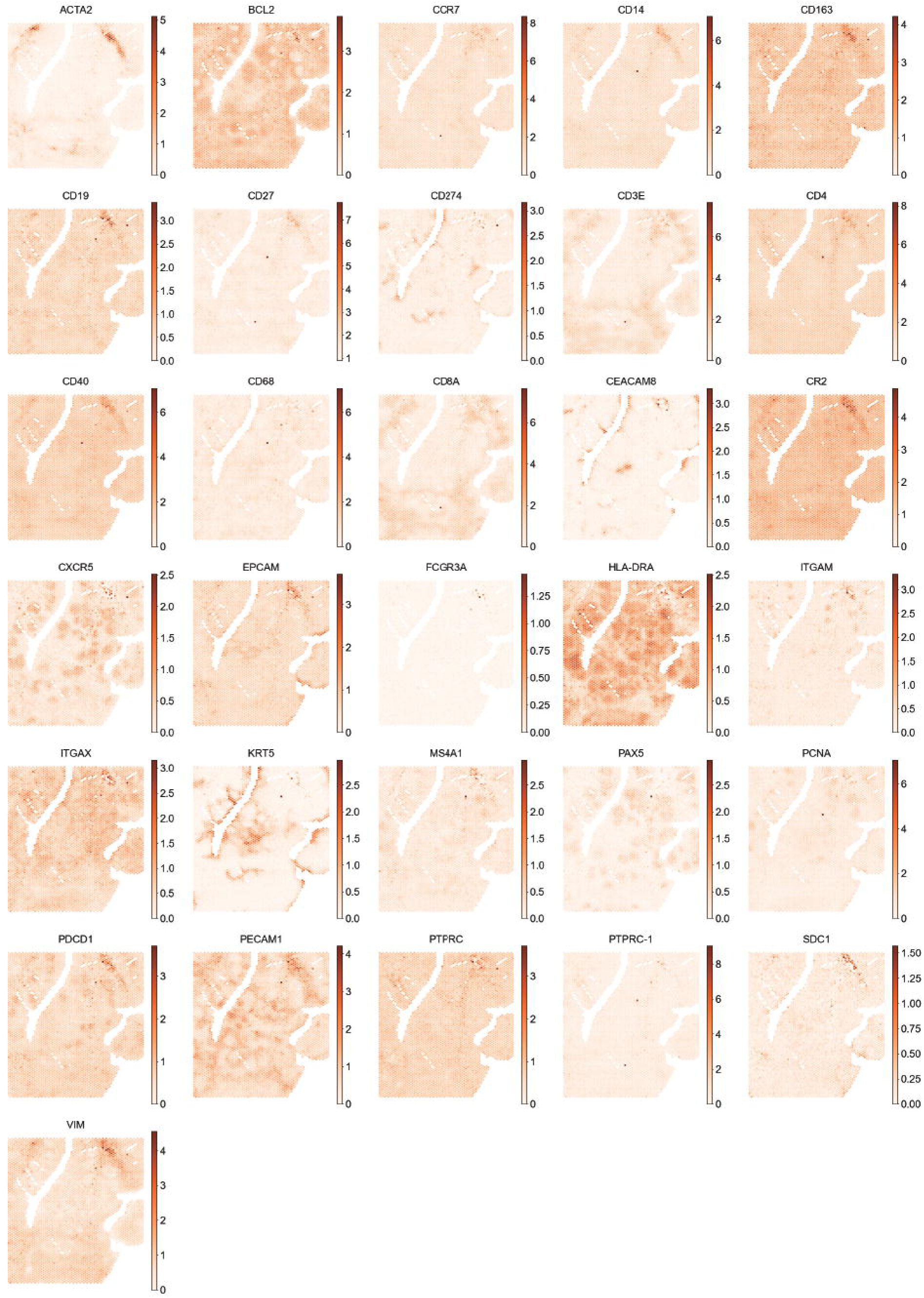
Spatial protein expression maps of all proteins measured in the CytAssist human tonsil section. Data are center-log-ratio normalized.

**Supp. Fig. S6.**
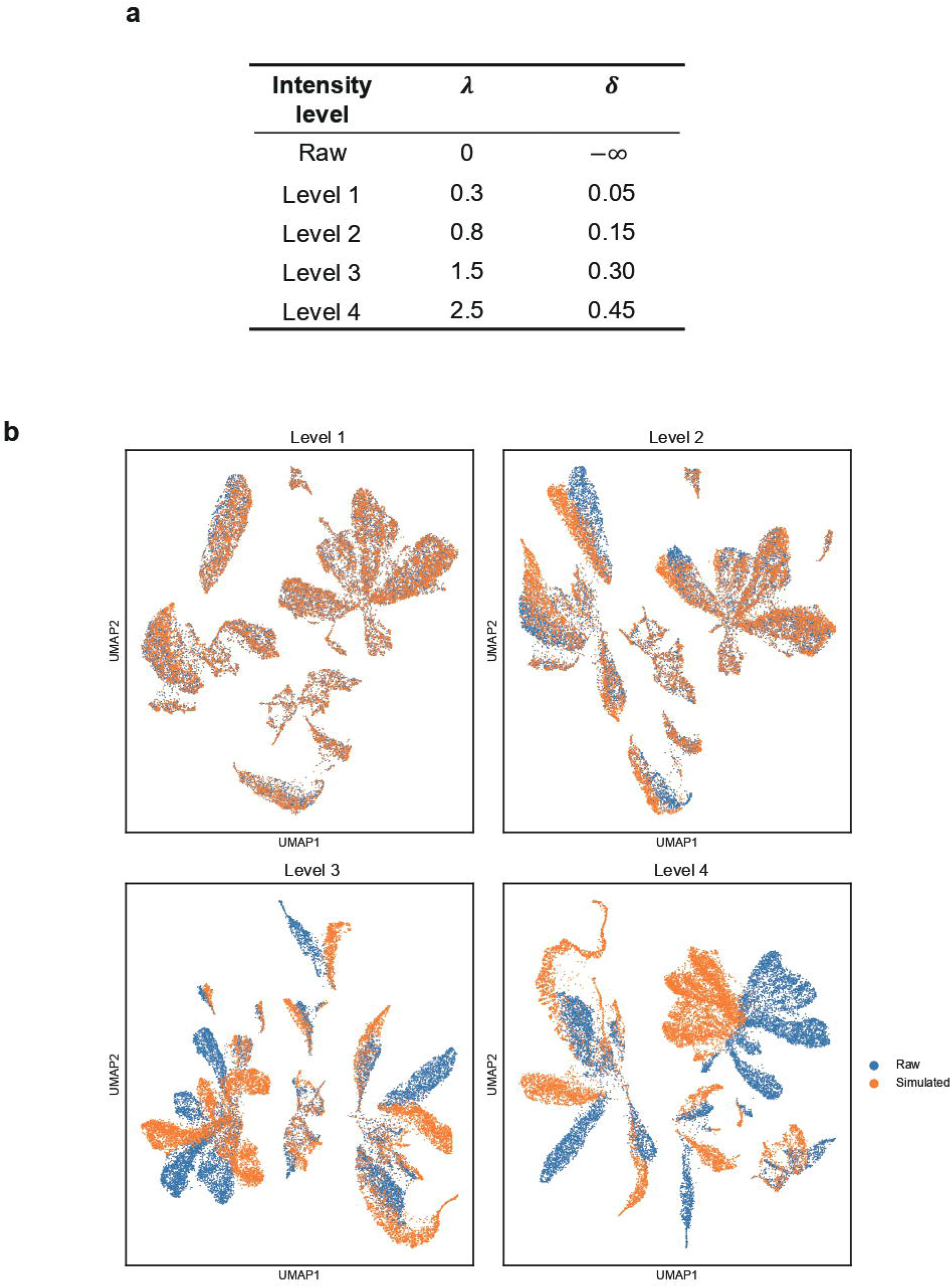
Reference perturbation parameters and visualization of their effects on an example mouse isocortex dataset. **a**, Perturbation parameters for the five intensity levels. **b**, UMAP visualization of the raw and perturbed reference data at each intensity level on an example dataset.

